# To let go or not to let go: how ParA can impact the release of the chromosomal anchoring in *Caulobacter crescentus*

**DOI:** 10.1101/2023.04.12.536610

**Authors:** Stephanie G. Puentes-Rodriguez, J.D. Norcross, Paola E. Mera

**Affiliations:** Department of Microbiology, University of Illinois at Urbana-Champaign, Urbana, IL, USA

## Abstract

Chromosomal maintenance is vital for the survival of bacteria. In *Caulobacter crescentus*, chromosome replication initiates at *ori* and segregation is delayed until the nearby centromere-like region *parS* is replicated. Our understanding of how this sequence of events is regulated remains limited. The segregation of *parS* has been shown to involve multiple steps including polar release from anchoring protein PopZ, slow movement, and fast ParA-dependent movement to opposite cell pole. In this study, we demonstrate that ParA’s competing attractions from PopZ and from DNA are critical for segregation of *parS*. Interfering with this balance of attractions – by expressing a variant ParA-R195E unable to bind DNA and thus favoring interactions exclusively between ParA-PopZ – results in cell death. Our data revealed that ParA-R195E’s sole interactions with PopZ obstruct PopZ’s ability to release the polar anchoring of *parS* resulting in cells with multiple *parS* loci fixed at one cell pole. We show that the inability to separate and segregate multiple *parS* loci from the pole is specifically dependent on the interaction between ParA and PopZ. Interfering with interactions between PopZ and the partitioning protein ParB, which is the interaction that anchors *parS* at the cell pole, does not rescue the ability of cells to separate the fixed *parS* loci when expressing *parA*-R195E. Thus, ParA and PopZ appear to have a distinct conversation from ParB yet can impact the release of ParB-*parS* from the anchoring at the cell pole. Collectively, our results reveal that the initial steps in chromosome segregation are highly regulated.

## INTRODUCTION

Chromosomal maintenance is vital for the survival of the cell. Unlike eukaryotes where the processes of chromosome replication and segregation are separate in time and space, bacteria replicate and segregate their chromosomes concurrently. Thus, the coordination between chromosome replication and segregation in bacteria must be tightly modulated to achieve proper chromosomal maintenance after the completion of each cell cycle. Any mistake in the coordination of these events can be lethal for the bacterial cell. The essentiality of this coordination makes these events prime targets for potential new antibiotics. However, our understanding of the complexities involved in the coordination of chromosome segregation post initiation of chromosome replication remains limited.

One of the most regulated steps during chromosome replication is the initiation, which is carried out by the canonical protein DnaA at the chromosomal locus referred to as *ori* (*ori*gin of replication) (1,2). DnaA is an AAA+ ATPase protein that oligomerizes at *ori* opening the double stranded DNA for the replication machinery (replisome) to initiate replication bidirectionally (3,4). The activity of DnaA is regulated at a multitude of levels highlighting the importance of tightly modulating this critical initial step (5). Furthermore, accumulating data from different bacterial species have exposed roles of DnaA that go beyond the initiation of chromosome replication. For instance, DnaA is also a transcription factor that can serve as a master regulator for the progression of the cell cycle (6–8). DnaA has been implicated in the coordination of other major cell cycle events like chromosome segregation and cytokinesis (9,10).

Unlike chromosome replication, the mechanisms that drive chromosome segregation appear to be more diverse among bacteria. The partitioning system ParABS has been shown to promote segregation of plasmids, chromosomes, and other types of large cytoplasmic cargos (11–17). Three components make up the ParABS system: two nucleotide-binding proteins (ParA & ParB) and one chromosomal locus known as the centromere-like region *parS*. ParB is a CTPase protein that binds *parS* and promotes the ATPase activity of ParA (18–20). The ParABS system is encoded in about 70% of bacterial species (21). When surveying the genomic location of this system in bacteria, *parS* was interestingly found to be located near *ori* (21–23). This genomic proximity between *ori* and *parS* supports the intrinsic connection between the initiation of chromosome replication and of chromosome segregation.

*Caulobacter crescentus* (referred here as *Caulobacter*) is an excellent model system where mechanistic details involved in the regulation of chromosomal dynamics have been identified (24). Although replication of the chromosome initiates at *ori*, chromosome segregation does not initiate until the centromere *parS* region (8kb away) is replicated (25). In non-replicating *Caulobacter* cells, *parS* is found anchored at the stalked pole by direct protein-protein interactions between ParB (bound to *parS*) and the anchoring protein PopZ (26,27). Three ordered steps have been proposed to be involved in the segregation of *parS* from the stalked pole to the opposite pole post replication initiation: 1^st^ *parS* is released from polar anchoring and initiates slow movement, 2^nd^ fast movement driven by ParABS, and 3^rd^ completion of *parS* segregation by anchoring with PopZ at the cell pole (25–30). The initial slow separation of the replicated *parS* loci has been proposed to be ParA-independent and driven by “bulk” segregation that includes various forces from DNA replication, transcription, and entropy forces (31,32). The asymmetric segregation of one of the replicated *parS* loci from the stalked pole to the opposite pole is maintained even in cells where the polarity of the chromosome organization was flipped, highlighting the robustness of the segregation machinery (33).

In this study, we explored the mechanisms involved in the initial separation of the replicated *parS* loci post chromosome replication initiation. We identified a variant of ParA that when expressed causes cells to retain their replicated *parS* loci fixed at the stalked pole inhibiting proper chromosome segregation. The apparent inability to separate the replicated *parS* loci is maintained even in cells that have multiple copies (>2) of *ori-parS* from over-initiation of chromosome replication. We show that the expression of this variant has a lethal effect that can be irreversible. This variant has a charge modification (R195E) in ParA’s DNA-binding domain preventing DNA binding, which makes this variant unable to trigger its ATPase activity (18,34,35). We discovered that the inability to separate *parS* in cells expressing *parA*-R195E is dependent on the chromosome anchoring protein PopZ. Removing PopZ or expressing a PopZ variant that is unable to interact with ParA rescues the ability to separate the replicated *parS* loci in cells expressing *parA*-R195E. Based on our results, we propose a model in which ParA’s function and localization is maintained by two attracting and competing forces coming from PopZ and DNA. Furthermore, our data suggest that the release of *parS* from the polar anchoring may be fine-tuned by other regulators making even the very first step in *parS* segregation more complex than previously thought.

## MATERIALS AND METHODS

### Strains and plasmids

All the strains and plasmids used in this study are listed in Supplementary Table 1 and 2. Plasmids construction was performed by cloning PCR-amplified DNA fragments into pXCHYC-2 and pVCHYC-6, (36) using *C. crescentus* wild-type CB15N (NA1000) genomic DNA as the DNA substrate. Plasmids were transformed into *E. coli* DH5α cells and grown in LB medium (Fisher Bioreagents) at 37°C with orbital shaking at 200 rpm. All plasmid constructs were sequenced to verify cloning.

### Growth conditions

*C. crescentus* strains were grown in minimal (M2G) or nutrient-rich media (PYE) from a freezer stock inoculum, at 30°C and 180 rpm. Liquid media was supplemented with 5 μg/mL Kanamycin (Kanamycin sulfate in water, IBI Scientific), 0.2 μg/mL Chloramphenicol (in 100% ethanol, Sigma Aldrich), 0.4 μg/mL Gentamycin (Gentamycin sulfate in water, Sigma Aldrich), 1.25 μg/mL Spectinomycin (Sigma Aldrich). PYE plates contained 25 μg/mL Kanamycin or 1 μg/mL Chloramphenicol. *C. crescentus* cells were synchronized using the mini-synchrony protocol to isolate a nascent population of swarmer cells (37). In short, cells were grown in M2G (15 mL) to OD600 ∼0.3 and pelleted by spinning at 6000 rpm for 10 min at 4°C. The cell pellet was resuspended in 800 μL of 1 × M2 salts, and then mixed with 900 mL Percoll (Sigma-Aldrich). The mixed solution was then transferred to a 2 mL microfuge tube and centrifuged at 11,000 rpm for 20 min at 4°C to separate the swarmer from stalked cells via a density gradient. The swarmer cells were extracted by collecting the bottom layer of the gradient, placed in another tube to be washed twice with 1 × M2 salts, centrifuging at 8000 rpm for 3 min at 4°C. The swarmer cells were resuspended in M2G medium (2 mL, OD600 ∼0.1). PYE medium was used during *C. crescentus* strain constructions.

### Microscopy

*C. crescentus* cells (∼2 μL) were spotted on agarose pads (1% agarose in M2G), then phase contrast fluorescence micrographs were obtained using the Zeiss Axio Observer 2.1 inverted microscope with AxioCam 506 mono camera (objective: Plan-Apochromat 100×/1.40 Oil Ph3 M27 [WD = 0.17 mm]), and Zen lite software. Analyses of data was performed with blinded samples. Number of foci per cell was manually counted (Cell Counter Plugin), and cell length was analyzed using ImageJ/FIJI with MicrobeJ software (38).

### Flow cytometry analysis

Swarmer cells were isolated via mini-synchrony (37) and resuspended into M2G medium. At different time points, cells were treated with a flow cytometry protocol (39) to analyze chromosome content. In short, cells were incubated with Rifampicin (16 μg/mL in 100% methanol) to halt DNA replication re-initiation, then incubated for 3 h at 30°C. Afterwards, the cells were fixed with 70% ethanol, and stored at 4°C for up to 24 h. Before flow cytometry analysis, the fixed cells were then spun down at 4000 × g and resuspended carefully in TMS buffer (10 mM Tris-HCl pH 7.2, 1.5 mM MgCl2, 150 mM NaCl), and stained with 10 μM Vybrant DyeCycle Orange (Invitrogen). The fixed samples were analyzed on a BD LSR II Flow cytometer, and the data were analyzed using BD FACSDiva software (BD Biosciences) and combined using Adobe Illustrator software.

### Genomic DNA analysis

*C. crescentus* cells were grown in M2G overnight at 30°C and 180 rpm, then induced for 1 hr with 0.1% xylose to promote the expression of different ParA variants. Then, the cells were synchronized to obtain a nascent population of swarmer cells, and 0.1% xylose was added for 2 hr. Afterwards, the genomic DNA content was extracted using the GeneJet Genomic DNA Purification Kit (Thermo Fisher) and sent for short-read sequencing (SeqCenter). Once the genomic reads were obtained, the data were analyzed using the Breseq pipeline to align chromosomal reads to the *Caulobacter* genome.

### Survival and Viability Assays

*C. crescentus* cells were grown in M2G from freezer stock inoculums, then re-inoculated in fresh media overnight at 30°C and 180 rpm. To perform Viability Assays, PYE agar plates were made with or without 0.1% xylose to induce the expression of different ParA variants, then the cultures were serially diluted and plated on the aforementioned plates. To perform Survival Assays, the cultures were induced with 0.1% xylose and incubated at 30°C and 180 rpm for 8 hr. Afterwards, the cultures were serially diluted and plated in PYE agar plates.

## RESULTS

### ParA’s inability to bind DNA results in trapped replicated ori-parS loci

We have previously shown that increased levels of ParA impact the regulation of chromosome replication initiation resulting in cells with multiple origins of replication (>2) in *Caulobacter* (40). In this study, we further our analyzes of ParA’s impact on chromosomal maintenance by focusing on the initial separation of the *ori-parS* loci post chromosome replication initiation. ParA-like proteins have DNA-binding domains that include several conserved positively charged amino acids lining both sides of the protein dimer interface (41). To analyze the role of ParA’s DNA-binding domain, we focused on one such conserved residue in *Caulobacter’*s ParA: Arg195 (41,42). We constructed merodiploid *Caulobacter* strains encoding the wildtype *parA* at its native promoter and the *parA*-(Arg195) variants at the xylose inducible promoter. To track chromosome replication and segregation, we fluorescently labeled the partitioning protein ParB, which binds the centromere-like locus *parS* near *ori* (25). Cells expressing a ParA variant with Arg195 replaced with Ala display multiple (>2) CFP-ParB foci (Figure 1A) suggesting that ParA’s ability to bind DNA is not required for its impact on chromosome replication initiation, consistent with our previous report (40). Because ParA-R195A is expected to retain reduced ability to bind DNA (43), we next explored the complete loss of DNA binding by replacing the positive charge of Arg with the negative charge of Glu (R195E). In *B. subtilis* Soj(ParA), the corresponding modification (R189E) was shown to eliminate the ability to bind DNA (43). The *Caulobacter* ParA-R195E variant has also been shown to lose the ability to bind DNA (35,42).

**Figure 1.**
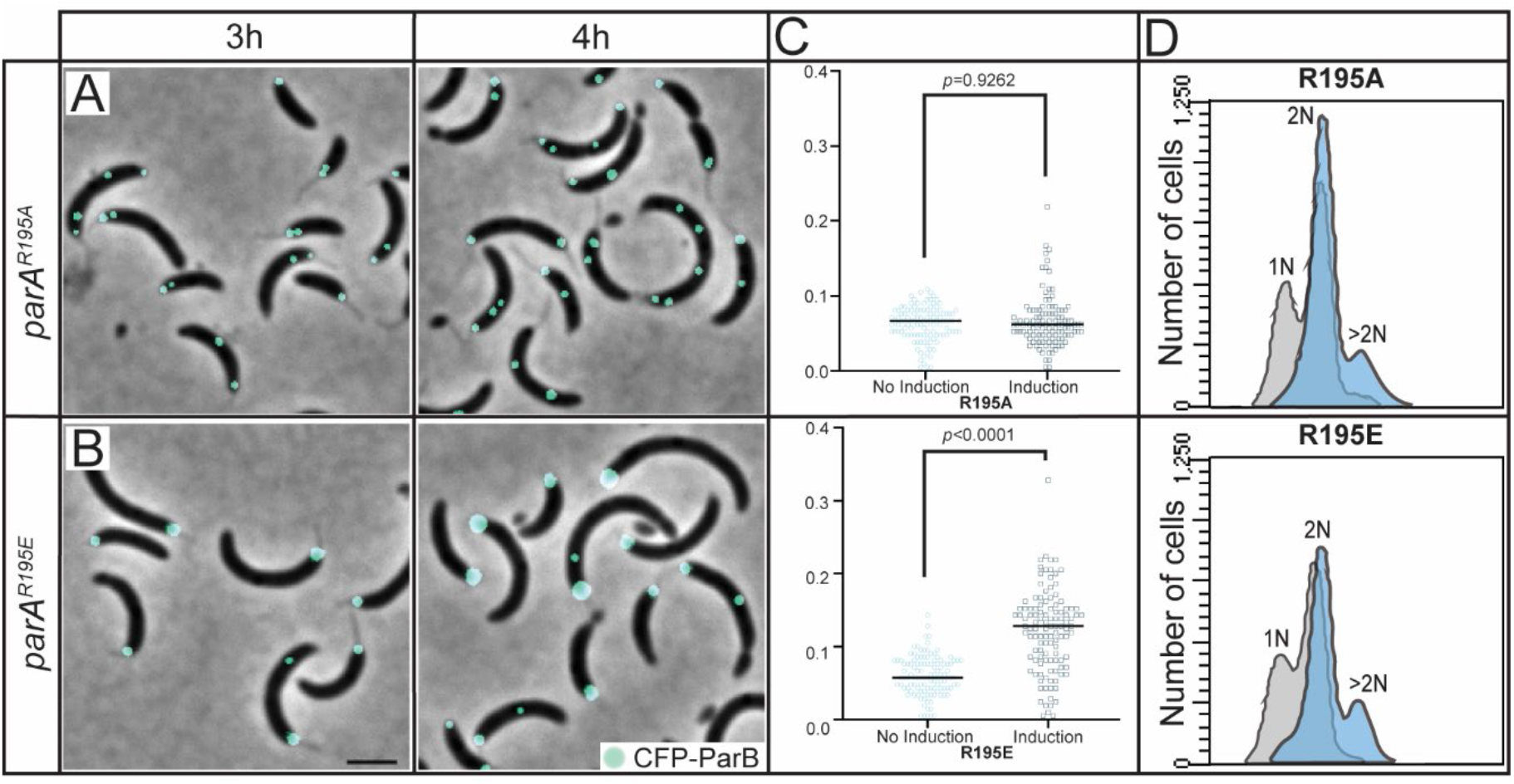
ParA^R195E^ disrupts the cell’s ability to separate replicated *parS* loci. A mixed population of CB15N cells expressing *parB::cfp-parB* (cyan) were diluted to 0.1 OD_600_, then added 0.1% xylose to express *xylX::parA-(R195A)* or *xylX::parA-(R195E)*, then micrographs were obtained at **A)** 3 h or **B)** 4 h of induction. Micrograph scale bar represents 2 µM. **C)** Flow cytometry profiles after 3 h of expression. Flow cytometry data was obtained by synchronizing the aforementioned cells and adding 0.1% xylose for 3 h, then adding Rifampicin to block re-initiation of chromosome replication for an additional 3 h. Chromosomal content is denoted above each peak. **D)** Analysis of ParB focus area in strains expressing *parB::cfp-parB and xylX::parA-(R195E)* or *xylX::parA-(R195A).* Cells were synchronized and added 0.1% xylose for 4 h, then micrographs were taken. Afterwards, analysis of ParB focus area was performed by counting foci area using FIJI, and statistical analyses were performed using two-way ANOVA, n=115.

Unlike from *parA*-R195A expression, cells expressing *parA*-R195E displayed a single CFP-ParB focus initially suggesting that this variant was unable to promote the over-initiation of chromosome replication (Figure 1B). However, the single focus of CFP-ParB in cells expressing *parA*-R195E increased in size overtime (Figure 1C & Supplemental Figure 1). This significant increase in foci size was not observed in the absence of induction of *parA*-R195E nor in cells expressing *parA*-R195A. To determine whether the larger CFP-ParB focus observed in cells expressing *parA*-R195E represented multiple copies of *parS*, we quantified chromosomal content per cell using flow cytometry. Our data revealed that cells expressing *parA*-R195E contain >2 chromosome equivalents (Figure 1D), just like cells that over-initiate replication from overexpression of wildtype *parA* (40) or in cells expressing *parA*-R195A. Collectively, these results revealed that cells expressing *parA*-R195E lose their ability to trigger the initial separation of replicated *parS* loci, even when cells contain more than 2 copies of *parS*.

### Expression of parA-R195E does not stop chromosome replication

Because changes in ParA levels can impact chromosome replication initiation in *Caulobacter* (40), we tested whether expression of *parA*-R195E could constrain the replication machinery. We used two strategies to address potential problems with chromosome replication in cells expressing *parA*-R195E. Firstly, we determined the localization of the replisome by using a strain encoding the β clamp (*dnaN*) fluorescently labeled at its native locus (44). Consistent with what has previously been shown in *Caulobacter* where the two replisomes work in concert (45), the majority of cells without induction of *parA* displayed a single focus or two side-by-side foci of DnaN-mCherry (Figure 2AB). On the other hand, cells expressing the variant ParA-R195A or ParA-R195E displayed multiple and separated DnaN-mCherry foci consistent with these cells undergoing multiple rounds of replication (Figure 1D & 2AB). Interestingly, under the expression of either variant (R195A or R195E), at least one DnaN-mCherry focus colocalized with a CFP-ParB focus. To determine whether this potential colocalization could impact chromosome replication, we analyzed the abundance of chromosomal regions by sequencing total DNA in cells expressing *parA*-R195E/A. Our goal was to test whether ParA-R195E could cause cells to replicate certain regions of the chromosome at higher frequency than others. Our data revealed that cells expressing *parA-*R195E display the same patterns of chromosomal abundance compared to cells expressing *parA*-R195A or wildtype (Figure 2C). Based on these results, we can posit that cells with CFP-ParB fixed at the stalked pole display no significant differences in the replication of the chromosome compared to cells with segregated *parS* foci. Thus, the inability to separate multiple CFP-ParB foci in cells expressing *parA*-R195E is not likely due to problems with chromosome replication.

**Figure 2.**
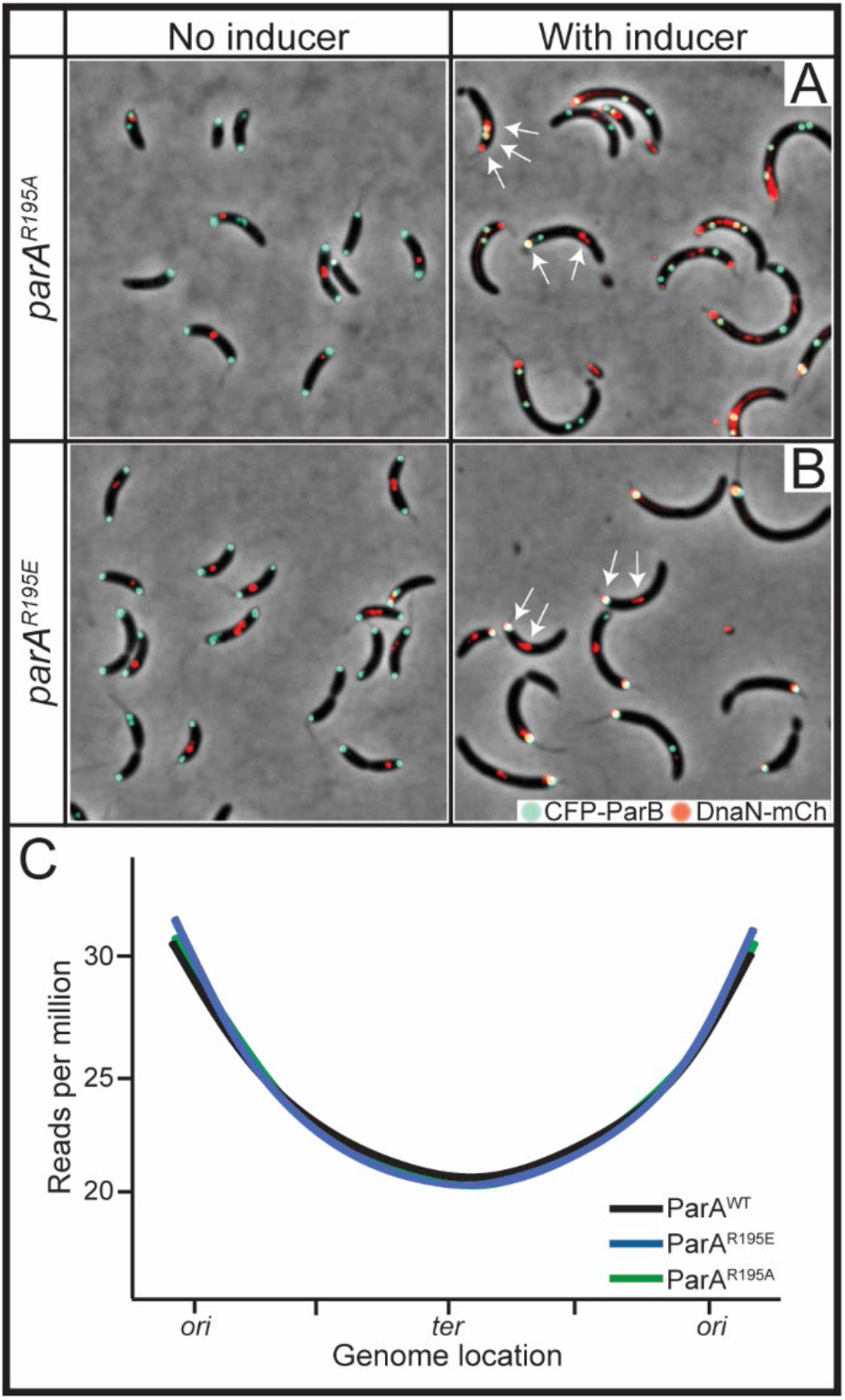
ParA^R195E^ does not inhibit progression of chromosome replication. CB15N cells expressing *parB::cfp-parB* and *dnaN::dnaN-mCherry* with **A)** *xylX::parA-(R195A)* or **B)** *xylX::parA-(R195E)* were diluted to 0.1 OD_600_, then added 0.1% xylose for 4 h. Arrows denote multiple DnaN-mCherry foci. **C)** Genomic DNA content of CB15N cells expressing *parB::cfp-parB* and either *xylX::parA-(WT), parA-(R195E)*, or *parA-(R195A)* for 2 h, with reads per million aligned to the genome location of *Caulobacter*. These cells were incubated with 0.1% xylose for 1 h prior to synchrony, then synchronized and incubated with 0.1% xylose for an additional 2 h before extracting the genomic content using a GeneJet Genomic DNA purification kit and sent to sequencing to SeqCenter. Data were analyzed using the Breseq pipeline to align reads to the *Caulobacter* genome.

### Chromosomal regions away from parS can separate when parA-R195E is expressed

Chromosomal regions near and distant (>100 kb) from *parS* segregate at different rates and are driven by different mechanisms (46,47). Because *ori* and *parS* are separated only by ∼8 kilobases (25), we tested the possible scenario where expression of *parA*-R195E also prevents the separation of the replicated *ori* loci. To test this hypothesis, we constructed a strain expressing *parA*-R195E with a chromosomal region near *ori* fluorescently labeled using the plasmid system (pMT1-*parS*-ParB) (48). Our data revealed that like *parS*, the replicated *ori* are also stuck at the cell pole and unable to separate, at least separate enough for fluorescent microscopy to distinguish two separated foci (Figure 3A). To confirm that the plasmid system pMT1(*parS*-ParB) was not interfering with the *Caulobacter* partitioning system, we used a separate strain with a fluorescent tag near *ori* using the *tetO*-TetR array system (49) and engineered the *parA*-R195E under the inducible promoter. Our data revealed that regardless of the system used, both strains expressing *parA*-R195E display *ori* loci as a single fluorescent focus (Figure 3B). Thus, cells expressing *parA*-R195E retain their *ori* and *parS* loci fixed at the stalked pole consistent with the close proximity of these two chromosomal loci (25). We next explored the impact of ParA-R195E on the organization of chromosomal loci distant from *parS*. To keep track of relative distances, we used a *Caulobacter* strain with two fluorescent labels: one at *parS* (native ParB fluorescently labeled) and one at 1.3 Mb from *parS* (pMT1-*parS*-ParB) near the gene *pleC* (50). Cells expressing *parA*-R195E with multiple *parS* loci stuck at the cell displayed separated foci of the chromosomal region near *pleC,* revealing that chromosomal regions distant to *parS* can separate in the presence of ParA-R195E (Figure 3C). Collectively, our analyses indicate that the expression of *parA*-R195E impacts mechanisms involved in the onset of chromosome segregation at or near *parS*, but not necessarily of chromosomal loci distant to *parS*.

**Figure 3.**
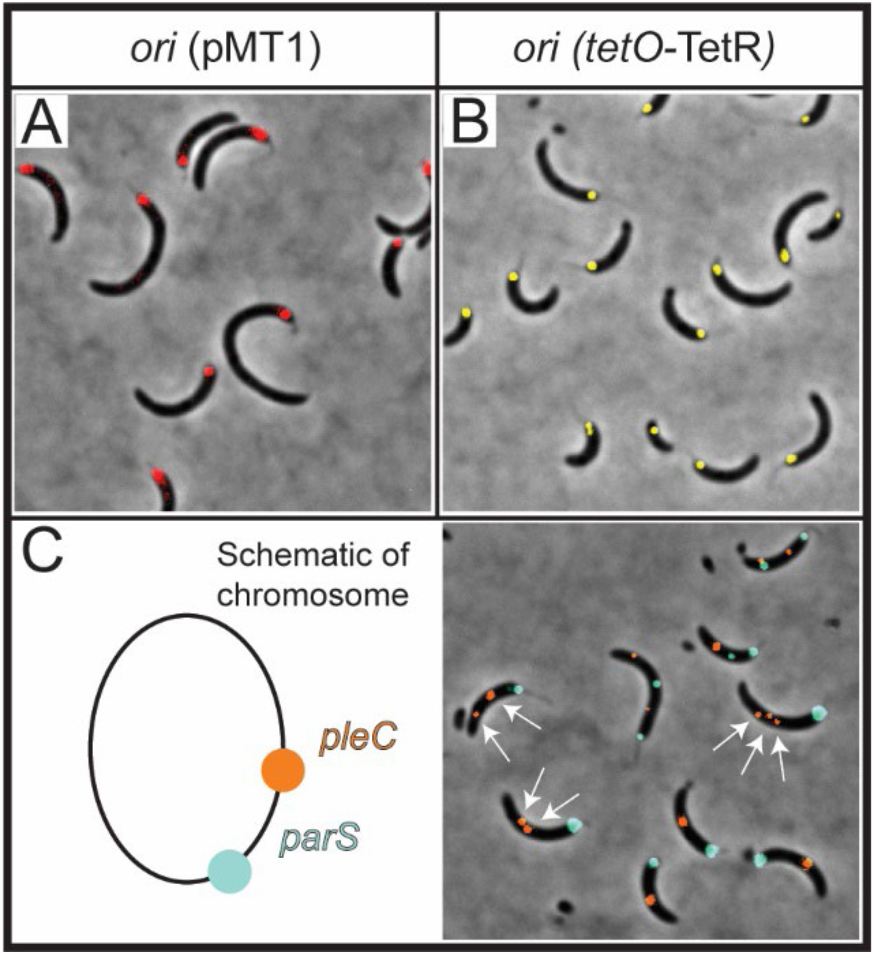
ParA^R195E^ impacts the separation of chromosomal loci near *parS*. **A)** Micrographs were obtained of a mixed population of CB15N cells expressing *parS (pMT1), xylX::yfp-parB* near *ori* (red) with 0.3% xylose and *vanA::parA-(R195E)* with 250 µM vanillate for 3 h. **B)** A mixed population of CB15N cells *with cc0006::(tetO)n, vanA::tetR-efyp* (yellow)*, xylX:: parA-(R195E)* with 0.1% xylose was observed, and micrographs were obtained after 4 h of induction. **C)** Micrographs of CB15N cells expressing *parB::cfp-parB* (cyan), *parS (pMT1)-pleC* at 1.3 Mb away from *ori, xylX::yfp-parB* (orange), and *vanA::parA-(R195E)* were obtained after 3 h of induction with 0.3% xylose and 250 µM vanillate. Cartoon depiction of the localization of *parS* (native) and *parS (pMT1)-pleC.* Micrographs are representative of the cell population.

### Expression of parA-R195E is lethal

We have demonstrated that increasing the levels of ParA wildtype and ParA variants that cannot bind DNA display different phenotypes. We next examined how these different phenotypes relate to cell viability. To determine the impact on viability in cells unable to separate their replicated *parS* loci, we performed colony-forming-unit (CFU) analyses. Exponentially growing cells were serially diluted and spotted on media plates containing xylose inducer for expression of wildtype or variants of *parA*. Increasing the levels of wildtype ParA displayed a loss in cell viability compared to our vector-control-only (Figure 4A). However, expression of either variant of ParA with defects in DNA binding resulted in cells almost completely dessimated. To determine whether the impact from expressing *parA*-R195E is reversible, we allowed cells to express *parA*-R195E and subsequently determined the ability of these cells to recover on media without inducer. Our data revealed that cells expressing *parA*-R195A or overexpressing WT *parA* were able to recover after 8 h induction (Figure 4B). However, cells expressing *parA*-R195E displayed a significant loss of viability compared to controls after the same time exposure. These results suggest that once cells have potentially trapped replicated *parS* at the stalked pole due to expression of *parA*-R195E, a significant number of cells are unable to reverse the effect after the expression of *parA*-R195E is removed.

**Figure 4.**
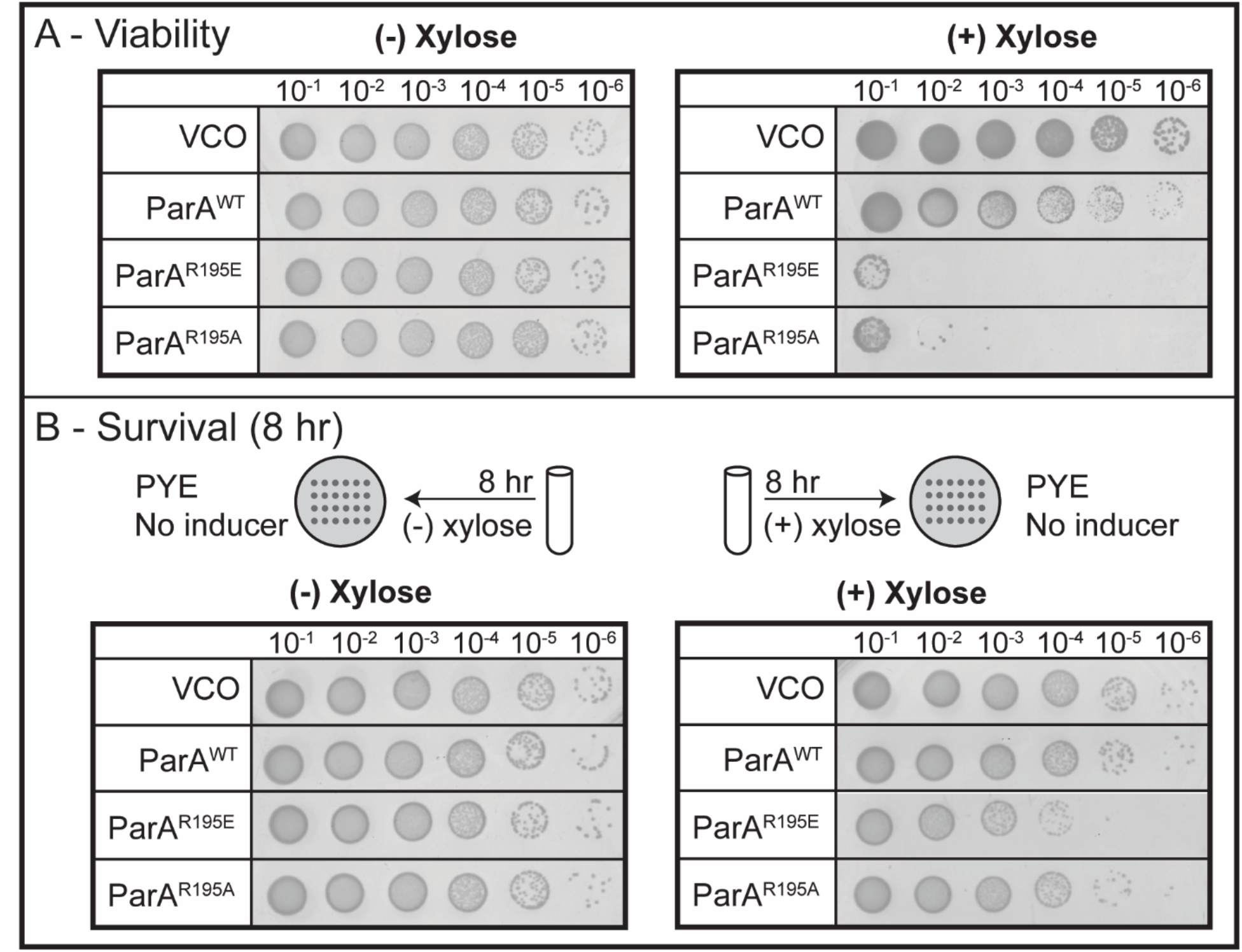
The expression of ParA^R195E^ is lethal to cells. **A)** Viability assays of CB15N cells with the genotype *parB::cfp-parB, and xylX::empty-vector*, *xylX::parA-(WT), xylX::parA-(R195E),* or *xylX::parA-(R195A)* were serially diluted before plating in PYE agar or PYE/0.1% xylose plates to induce the expression of the aforementioned genes under xylose. Afterwards, plates were incubated at 30°C for two days before imaging. **B)** Survival Assays of the aforementioned cells incubated with 0.1% xylose for 8 h in M2G medium, then serially diluted to be plated in PYE agar plates and incubated for two days at 30°C. Data are representative of triplicate assays.

### Loss of DNA-binding impacts ParA’s cellular localization

The ability of ParA to dynamically interact with the chromosome serves to establish ParA’s cellular gradient localization (51). Our data revealed that neither variant ParA-R195E nor ParA-R195A can form the gradient structure observed with wildtype ParA and instead both form foci at the poles (Figure 5AB). Notably, we observed two differences in the localization between ParA-R195E and ParA-R195A that provided clues to why one ParA variant could separate the replicated *parS* loci but not the other. Unlike ParA-R195A, ParA-R195E exclusively localizes at the stalked pole and colocalizes with ParB (Figure 5BDE). The interesting difference between these ParA variant proteins is their localization in reference to the cell pole. A required step in the ordered multistep process involved in *parS* segregation in *Caulobacter* is the release of *parS*-ParB from the cell pole (52). Consistent with this requirement, we observed a sub-population of cells expressing *parA-*R195A display the CFP-ParB focus localized slightly away from the stalked pole (Figure 5AC). Our ability to capture these subtle changes in CFP-ParB localization was facilitated by ParA-R195A-mCherry foci which localized closer to the cell pole than CFP-ParB. However, we did not observe cells expressing *parA*-R195E display CFP-ParB localized away from the cell pole. The lack of separation of *parS*-ParB from the stalked pole in cells expressing *parA*-R195E exposed a potential problem with the anchoring of *parS*-ParB at the cell pole

**Figure 5.**
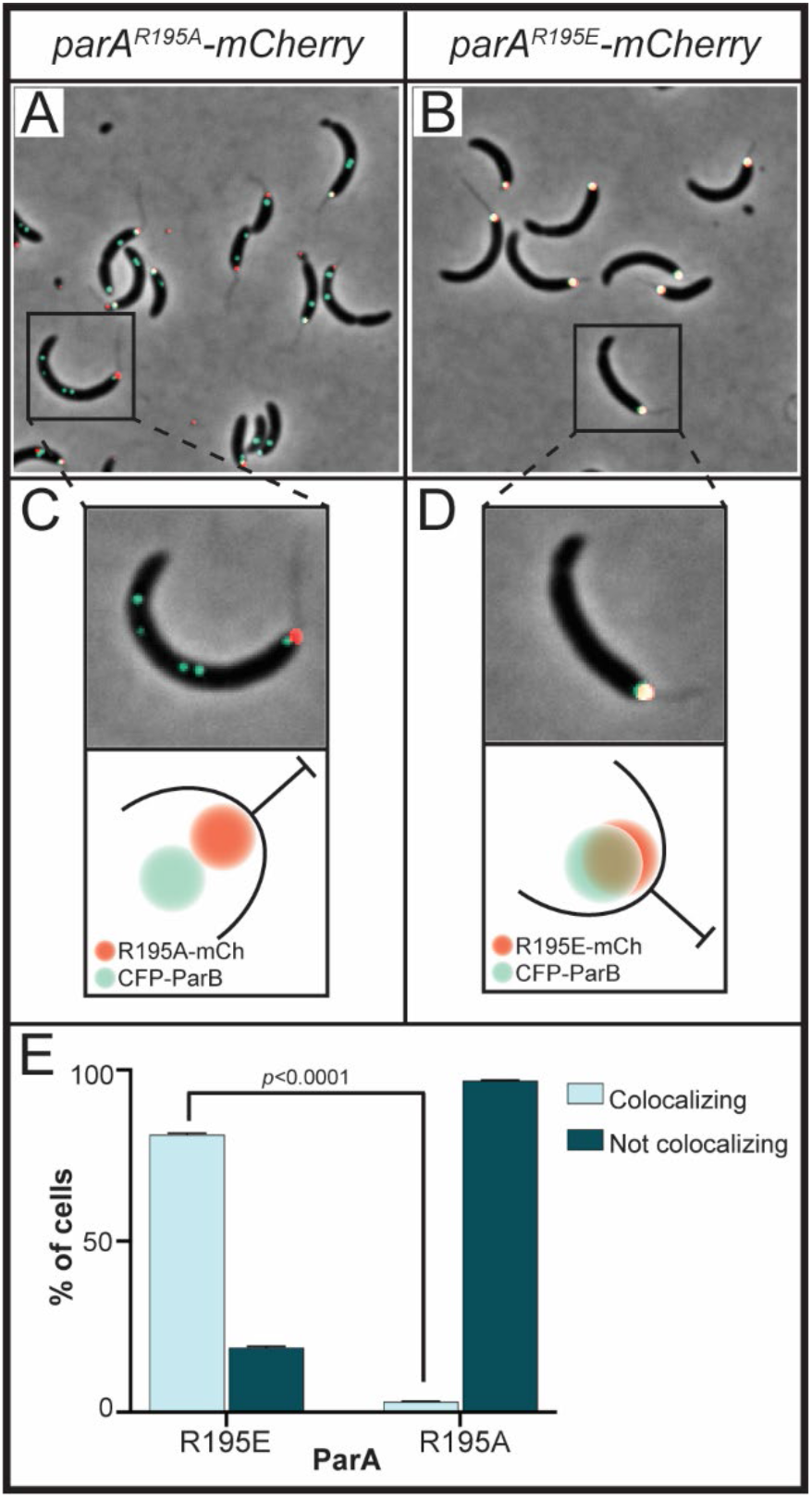
ParA^R195E^ colocalizes with *parS-*ParB at the stalked cell. Micrographs of CB15N cells expressing *parB::cfp-parB* and **A)** *xylX::parA-(R195A)-mCherry* or **B)** *xylX::parA-(R195E)-mCherry* were obtained by diluting a mixed population to 0.1 OD_600_, then incubating with 0.1% xylose for 4 h before imaging. **C, D)** Close-ups and a cartoon depiction of the localization of CFP-ParB and ParA-mCherry at the stalked cell pole. **E)** Quantification of colocalization of CFP-ParB with ParA-mCherry. Data represent error bars of mean ± SD and were blindly counted using ImageJ and analyzed using two-way ANOVA statistical analyses.

### Monopolar localization of ParA-R195E requires the anchoring protein PopZ

#### (53)-popZ and microdomain

*Caulobacter* cells anchor their chromosome at the stalked pole by direct interactions between ParB (bound to *parS*) and the anchoring protein PopZ (54,55). Based on our observation that *parS*-ParB remain fixed at the stalked pole in cells expressing *parA*-R195E, we hypothesized that ParA-R195E could disturb the anchoring function of PopZ. To test this hypothesis, we first analyzed the cellular localization of PopZ using a strain with the native *popZ* fluorescently labeled (54) as the background for the xylose-inducible expression of *parA* variants. In *Caulobacter*, PopZ localizes at one cell pole (stalked pole) in non-replicating cells and at both cell poles in actively growing cells (54,55). We tracked the localization of PopZ as the expression of *parA*-R195E/A were induced over time. We used strains with ParA variants fluorescently labeled to correlate impact on PopZ’s localization with appearance of ParA variants. Our time-course analyzes revealed that upon 3 h of induction, most cells display fluorescent foci for the ParA variants (Supplementary Figure 2). Based on these results, we analyzed the percent of cells displaying monopolar and bipolar PopZ localization (Figure 6AB). When *parA-* R195A is expressed, cells display one or two PopZ-YFP foci localized at the cell poles, just like what is observed in the control cells with empty vector. These data revealed that cells expressing *parA-*R195A retain the ability to bipolarly localize PopZ. However, cells expressing *parA*-R195E are no longer able to bipolarly localize PopZ (Figure 6). In the presence of ParA-R195E, ∼70% of cells displayed only one PopZ focus, which exclusively localized at the stalked pole. We conclude from these data that expression of *parA*-R195E impacts the ability of cells to maintain the proper localization of PopZ.

**Figure 6.**
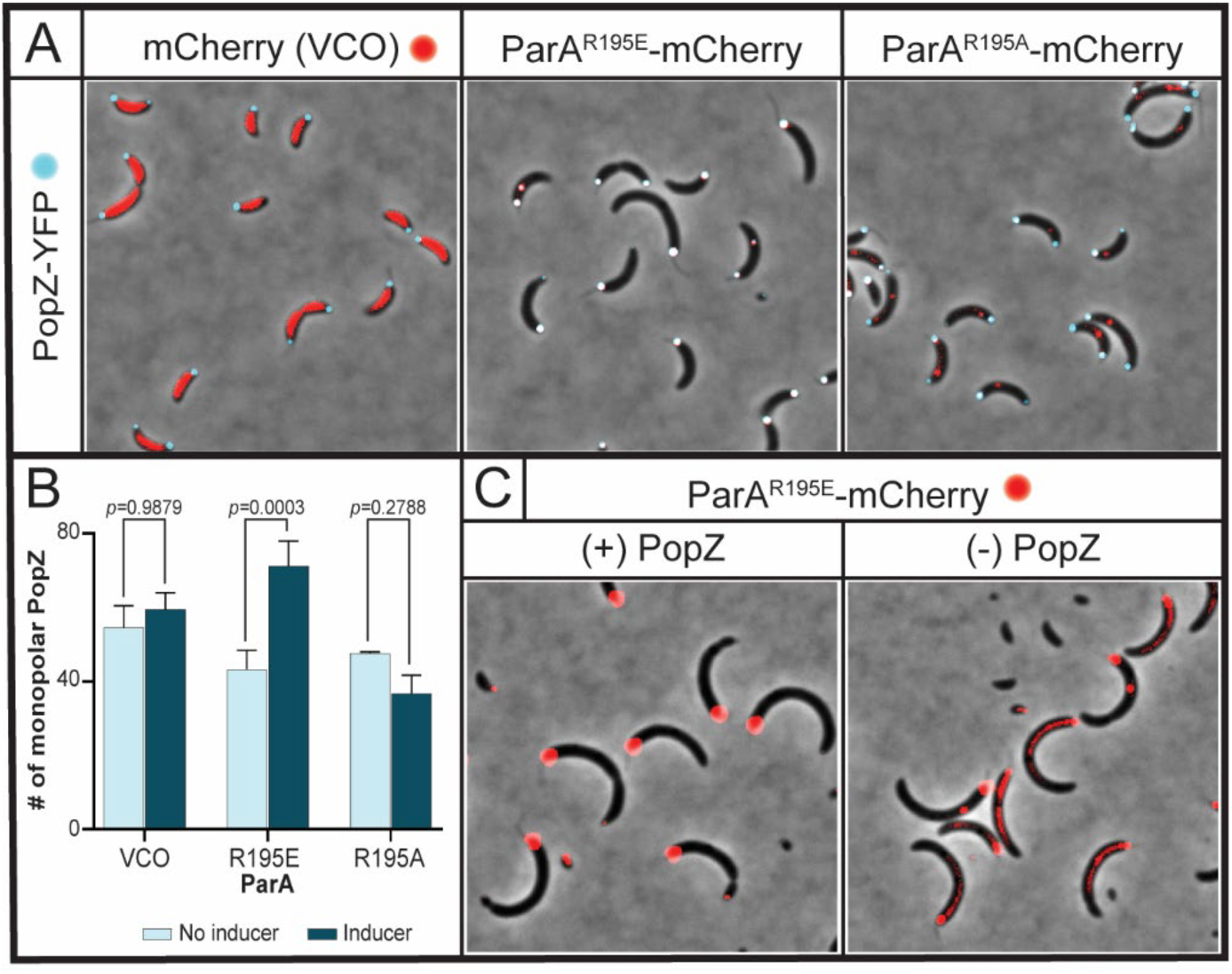
Localization of ParA^R195E^ at the stalked pole requires PopZ. Micrographs of CB15N cells expressing *popZ::popZ-YFP* and **A)** *xylX::mCherry*, *xylX::parA-(R195E)-mCherry*, or *xylX::parA-(R195A)-mCherry*. Images were obtained by diluting a mixed population to 0.1 OD_600_, then incubating with 0.1% xylose for 3 h. **B)** Quantification of monopolar PopZ foci in cells expressing *xylX::mCherry*, *xylX::parA-(R195E)-mCherry*, or *xylX::parA-(R195A)-mCherry*. Data was blindly counted using ImageJ and analyzed using two-way ANOVA statistical analyses. Bar graphs represent error bars of mean ± SD from three independent replicates. **C)** A mixed population of CB15N cells expressing *xylX::parA-(R195E)-mCherry* and *vanA::popZ* were grown overnight in M2G with 250 µM vanillate, then washed with 1x M2 salts to remove vanillate. Afterwards, the cells were incubated for 4 h with (left) or without (right) 250 µM vanillate before micrographs were obtained. Images are representative of the cell population.

In the vector-control-only, cells expressing mCherry (with no *parA*) display bipolar localization of PopZ with no overlap with mCherry (Figure 6A). In other words, the mCherry protein does not enter the PopZ microdomain at the cell poles consistent with PopZ’s strict specificity (53,56). Unlike mCherry alone, ParA-R195E-mCherry shows colocalization with PopZ at the stalked pole. To determine whether PopZ is required for the localization of ParA-R195E at the cell pole, we analyzed a PopZ depletion strain (54) with the inducible expression of *parA*-R195E. In the presence of PopZ, ParA-R195E-mCherry also localizes at the stalked pole (like PopZ) displaying a large fluorescent focus (Figure 6C). Notably, when PopZ is depleted, the localization of ParA-R195E-mCherry transitions from polar to cytoplasmic. Collectively, these data suggest that PopZ may be involved in recruiting ParA-R195E to the stalked pole.

### The role of the anchoring protein PopZ in the separation of replicated parS

Our data has so far revealed that cells expressing *parA*-R195E retain the replicated *parS* loci fixed at the stalked pole where ParA-R195E and PopZ localize. Based on these results, we hypothesized that ParA-R195E could prevent PopZ from releasing the anchoring of *parS*-ParB from the stalked pole for chromosome segregation to initiate. If the hypothesis were correct, we reasoned that eliminating ParA-R195E’s localization at the stalked pole would result in cells able to separate replicated *parS* loci. To test this hypothesis, we constructed a strain with *parA*-R195E’s expression regulated under the xylose promoter, and the fluorescently labeled ParB regulated under its own promoter with the *popZ* knockout strain as the background. Unlike cells with *popZ*, our data revealed that PopZ depletion and expression of *parA*-R195E results in cells displaying multiple and importantly separated *parS*-ParB foci (Figure 7A). In other words, cells expressing *parA*-R195E are able to separate the replicated *parS* loci as long as ParA-R195E does not localize at the stalked pole.

**Figure 7.**
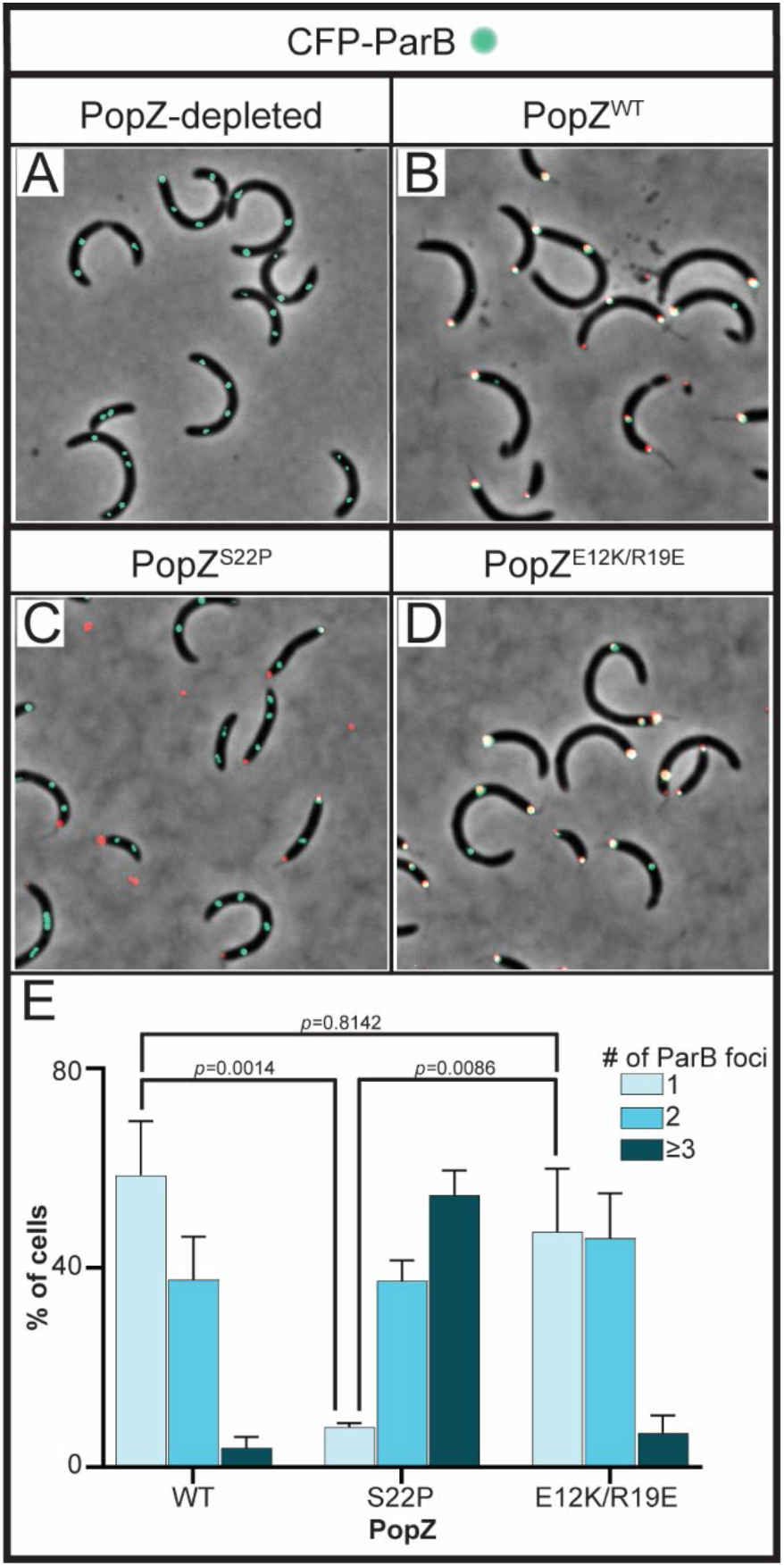
Disruption of ParA-PopZ interactions restores cells’ ability to separate replicated *parS* loci. Micrographs of CB15N cells expressing *parB::cfp-parB* (cyan), *xylX::parA-(R195E)* and **A)** *popZ::mCherry-popZ-(WT)*, **B)** *popZ::mCherry-popZ-(S22P)*, or **C)** *popZ::mCherry-popZ-(E1K2K/R19E)* (red). **D)** Quantification of ParB foci in the aforementioned cells. Data represent error bars of mean ± SD and analyzed with two-way ANOVA statistical analyses.

Because PopZ is known to house several key regulators that diffuse in and out of the PopZ microdomain at the cell poles (54,56), we further examined whether the ParA-R195E’s effect on the separation of replicated *parS* was independent from PopZ’s ability to form polar microdomains. To do that, we analyzed a fluorescently labeled PopZ variant with a single amino acid modification (PopZ-S22P) that retains its ability to form polar microdomains but is unable to interact specifically with ParA (57). Excitingly, our data revealed that cells expressing *parA*-R195E and *popZ*-S22P (from native promoter) displayed multiple and separated *parS* loci (Figure 7BC), similarly to the PopZ depletion strain (Figure 7A). In fact, cells expressing *popZ*-S22P were indistinguishable when grown in the presence or absence of the inducer for *parA*-R195E expression (Supplementary Figure 3). These data revealed that once ParA-R195E cannot interact with PopZ it no longer impacts the release of the replicated *parS* loci even when the microdomain of PopZ is formed at the cell poles.

Because PopZ directly interacts with ParB and this interaction is crucial for the release of the anchoring of *parS*, we examined whether the ParA-R195E’s effect on the separation of replicated *parS* loci was independent from PopZ’s interaction with ParB. The PopZ-S22P variant unable to interact with ParA was shown to retain its ability to interact with ParB (57) suggesting that the effect we observed was specific to the interaction with ParA. To further test this hypothesis, we analyzed the variant PopZ-E12K/R19E which was shown to lose the ability to interact with ParB and retain the ability to interact with ParA (57). Remarkably, cells expressing *popZ*(E12K/R19E) unable to interact with ParB displayed the single CFP-ParB focus observed in cells expressing wildtype PopZ (Figure 7D). These data revealed that ParA-R195E’s impact on PopZ’s ability to release *parS*-ParB is independent from PopZ’s interaction with ParB. Collectively, our work demonstrates that PopZ, besides DNA, is a key player in the regulation of ParA’s localization and function.

## DISCUSSION

In this study, we uncovered the impact that destabilizing the DNA-ParA interactions can have on the release of the chromosomal anchoring at the cell poles. Our data revealed that expression of a ParA variant that is unable to bind DNA results in cells unable to separate the replicated *parS* chromosomal loci. We found this inability to separate to persist even in cells over-initiating chromosome replication containing multiple (>2) *ori-parS* loci. Our data confirmed that the cell’s ability to separate the *ori-parS* regions post-initiation of chromosome replication is essential for viability. When the anchoring protein PopZ was removed or mutated to prevent specific PopZ-ParA interactions, cells were then able to release the replicated *parS* loci from the polar anchoring. Collectively, our data suggest a model in which PopZ-ParA interactions are critical for maintaining the proper localization of ParA not only for successful chromosome segregation but also for the initial separation of *ori-parS* loci post chromosome replication initiation (Figure 8).

**Figure 8.**
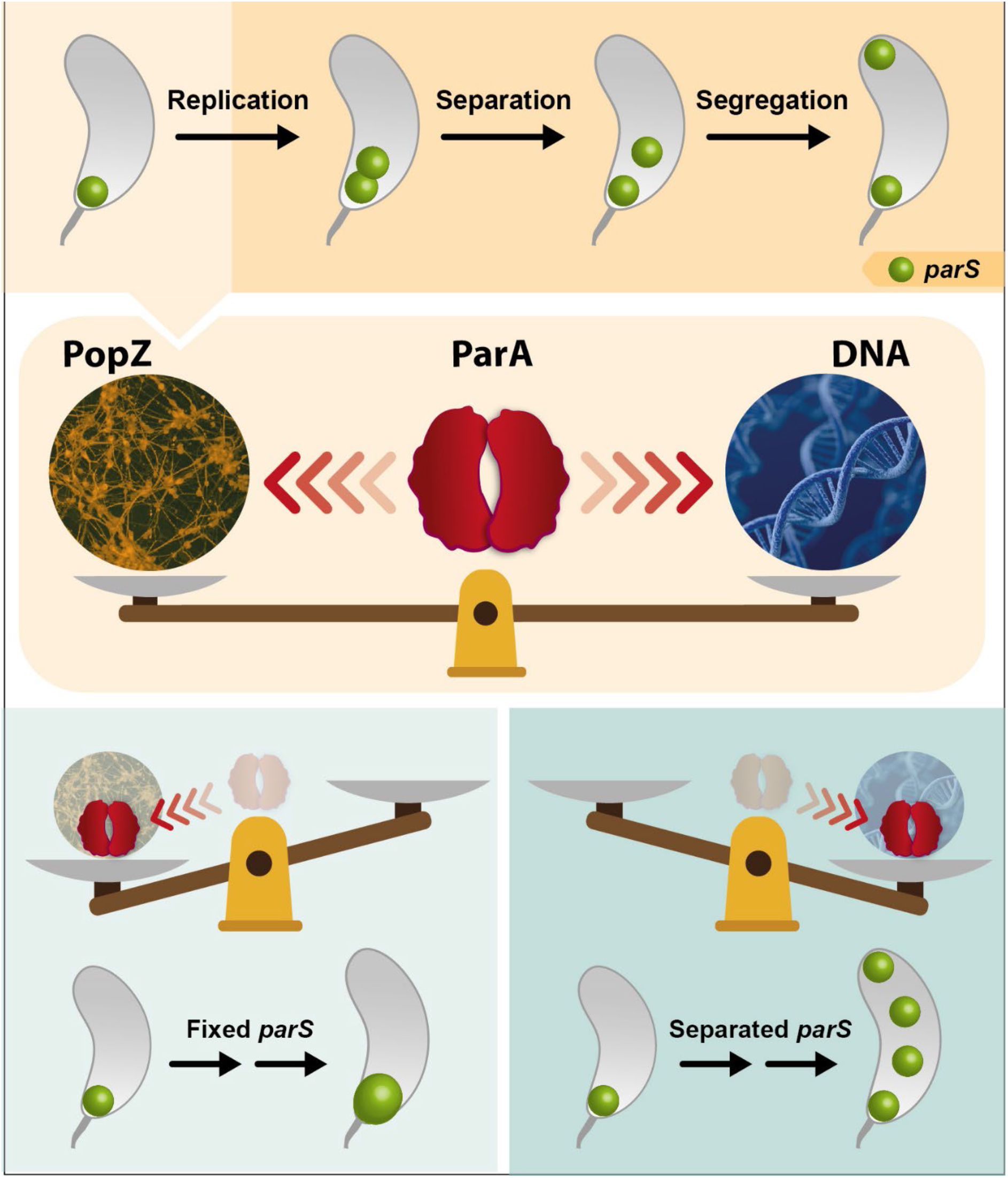
Summary Figure. Segregation of the centromere-like region *parS* involves the coordination of multiple events. This coordination of events requires the maintenance of ParA’s localization for proper completion of parS segregation. The localization of ParA is maintained by balancing between two attracting forces coming from PopZ and from the chromosome. When this balance is perturbed by favoring either PopZ or DNA, cells lose their ability to coordinate the replication and segregation of the chromosome.

We have shown that ParA’s inability to bind DNA results in the replicated *parS* loci to remain fixed at the cell pole. Previous analyses *in vitro* have shown that ParA-R195E’s inability to bind DNA also results in its inability to hydrolyze ATP (58). Therefore, the question that arises is whether the lack of ATPase activity of ParA-R195E could cause the *parS*-ParB foci to get stuck at the cell pole. We do not favor this hypothesis because we have previously shown that cells expressing a ParA variant (D44A) that can bind DNA but cannot hydrolyze ATP do not display the multiple *parS*-ParB foci stuck at the cell pole (40). In fact, the expression of *parA*-D44A results in cells displaying the same frequency of multiple (>2) separated *parS*-ParB foci as cells with overexpression of wildtype *parA*. Based on these data, we conclude that the inability of ParA-R195E to bind DNA, and not its lack of ATPase activity, results in cells unable to separate *parS* loci post chromosome replication initiation.

What form of ParA does PopZ interact with? PopZ has been shown to interact with various oligomeric states of ParA. Using super-resolution microscopy, cells overexpressing *popZ* with enlarged PopZ microdomains were shown to colocalize with the monomer of ParA-ATP (57). Using *E. coli*, which lacks ParABS, as a reporter system, dimers of ParA were shown to colocalize with PopZ at the cell pole (35). Using *in vitro* analyzes (surface plasmon resonance), PopZ was shown to favor direct interactions with ParA+ATP (dimer) and low interactions with ParA in the absence of nucleotide (57). Our data revealed that dimers of ParA, unable to bind DNA, colocalize with PopZ. Collectively, these data suggest that different forms of ParA may be able to interact with PopZ. One possibility is that PopZ interacts with both monomers and dimers of ParA because PopZ can itself promote changes in the nucleotide state of ParA and thus serve a more involved role in chromosome segregation. This model would be consistent with previous work showing that disrupting ParA-PopZ interaction can result in more severe defects than disrupting PopZ-ParB interactions (57). The possibility of PopZ modulating ParA’s activity is an exciting hypothesis that needs further analyzis.

ParA’s intrinsic low ATPase-independent rate of binding and unbinding to DNA was previously demonstrated to be critical for establishing the ParA’s gradient localization prior to triggering chromosome segregation (51). Our work adds to this model the role of PopZ’s interaction with ParA to maintain ParA’s proper localization and generation of its gradient. Our data revealed that impacting the balance between PopZ and DNA results in loss of ParA gradient localization. Cells expressing either variant (R195E or R195A) were unable to form gradients and instead localized at the cell poles, with ParA-R195E only found at the stalked pole. We propose that *Caulobacter* has optimized ParA’s affinity for PopZ and DNA to achieve a critical balance that allows for proper chromosome segregation. If the balance is disturbed in either direction by expressing *parA*-R195E (favoring the PopZ side) or expressing *popZ*-S22P with wildtype *parA* (favoring the DNA side), cells lose their ability to regulate chromosome replication and segregation (Figure 8). This is consistent with previous observations where expression of *parA*-R195E in cells with enlarged PopZ microdomains (caused by overexpression of *popZ*) also display single enlarged ParB foci at the stalked pole (59). Notably, in cells with enlarged PopZ microdomains, the release of *parS*-ParB foci was also accomplished when the PopZ variant missing its N-terminus was expressed instead of wildtype *popZ*. The N-terminus domain of PopZ interacts with ParA and ParB (57,59,60). Our data suggest that PopZ’s ability to let go of the anchoring of *parS*-ParB could be modulated by ParA independently of the direct interaction between PopZ and ParB. More work is necessary to unveil critical details of the complex conversations happening between the partitioning system and the anchoring protein PopZ for chromosome segregation.

The role of ParA with chromosomal maintenance in bacteria is turning out to be more complex than originally expected. Even though mechanistic details involved in ParA function varies among bacterial species, the extensive roles of ParA with the cell cycle appear to be conserved. For instance, the *B. subtilis* ParA(Soj)’s affinity for DNA is specific for the *ori* region whereas *Caulobacter* ParA’s affinity is nonspecific (25,35,42,61). ParA(Soj) has been shown in various bacterial species to impact directly or indirectly the rates of chromosome replication initiation (40,61,62). Recent work in *B. subtilis* has shown that ParA(Soj) and its various oligomeric nucleotide-dependent states can serve additional roles (63).

One of the initial steps during sporulation in *B. subtilis* is the reorganization of the chromosome in a structure known as the axial filament (64). The formation of the axial filament requires the anchoring of the chromosome at the cell poles by polar proteins like DivIVA and RacA (65–67). Consistent with our observation of ParA able to impact centromere anchoring, the monomer of ParA(Soj) was shown to impact the formation of the axial filament during sporulation (63). Thus, conversations between polar proteins and ParA-type partitioning proteins may be a widely used strategy to drive chromosomal maintenance in bacteria.

## ACKNOWLEDGEMENTS

We are very appreciative of various laboratories who generously shared strains and plasmids with us including Dr. Lucy Shapiro from Stanford University, Dr. Grant Bowman from the University of Wyoming, and Dr. Peter Chien from the University of Massachusetts. We thank Berent Aldikacti (Chien’s Lab) and David Vereau Gorbitz for assistance with the analyses of genomic abundance data, and Mr. Israel Ponce for the construction of the summary figure.

## FUNDING INFORMATION

The work reported in this publication was supported by the National Institute of General Medical Sciences of the National Institutes of Health NIH under [Award Number R01GM133833 to PEM]. S.G. P-R. was supported by R01GM133833-Supplement.

## CONFLICT OF INTEREST DISCLOSURE

The authors declare no conflict of interests.

## DATA AVILABILITY

All data are contained within the manuscript.

## Notes

### Competing Interest Statement

The authors have declared no competing interest.

## REFERENCES

1. Messer, W. (2002) The bacterial replication initiator DnaA. DnaA and oriC, the bacterial mode to initiate DNA replication. FEMS Microbiology Reviews 26, 355–374.

2. Boye, E., Lobner-Olesen, A. and Skarstad, K. (1988) Timing of chromosomal replication in Escherichia coli. Biochimica et Biophysica Acta, 951, 359–364.

3. Bramhill, D. and Kornberg, A. (1988) Duplex Opening by dnaA Protein at Novel Sequences in Initiation of Replication at the Origin of the *E. coli* Chromosome. Cell, 52, 743–755.

4. Marszalek, J. and Kaguni, J.M. (1994) DnaA Protein Directs the Binding of DnaB Protein in Initiation of DNA Replication in *Escherichia coli*. The Journal of Biological Chemistry, 269, 4883–4890.

5. Katayama, T., Ozaki, S., Keyamura, K. and Fujimitsu, K. (2010) Regulation of the replication cycle: conserved and diverse regulatory systems for DnaA and oriC. Nature Reviews Microbiology, 8.

6. Messer, W. and Weigel, C. (1997) DnaA initiator—also a transcription factor. Molecular Microbiology, 24, 1–6.

7. Hottes, A.K., Shapiro, L. and McAdams, H.H. (2005) DnaA coordinates replication initiation and cell cycle transcription in Caulobacter crescentus. Molecular Microbiology, 58, 1340–1353.

8. Menikpurage, I.P., Woo, K. and Mera, P.E. (2021) Transcriptional Activity of the Bacterial Replication Initiator DnaA. Front Microbiol, 12, 662317.

9. Mera, P.E., Kalogeraki, V.S. and Shapiro, L. (2014) Replication initiator DnaA binds at the Caulobacter centromere and enables chromosome segregation. Proceedings of the National Academy of Sciences, 111, 16100–16105.

10. Scholefield, G., Veening, J.-W. and Murray, H. (2011) DnaA and ORC: more than DNA replication initiators. Trends in Cell Biology, 21.

11. Abeles, A.L., Friedman, S.A. and Austin, S.J. (1985) Partition of Unit-copy Miniplasmids to Daughter Cells III. The DNA Sequence and Functional Organization of the Pl Partition Region journal of Molecular Biology, 185, 261–472.

12. Austin, S. and Abele, A. (1983) Partition of Unit-copy Miniplasmids to Daughter Cells II. The Partition Region of Miniplasmid P1 Encodes an Essential Protein and a Centromere-like Site at which it Acts. Journal of Molecular Biology, 169, 373–387.

13. Austin, S. and Abele, A. (1983) Partition of Unit-copy Miniplasmids to Daughter Cells I. P1 and F Miniplasmids Contain Discrete, Interchangeable Sequences Sufficient to Promote Equipartition. Journal of Molecular Biology, 169, 353–372.

14. Mori, H., Kondo, A., Ohshima, A., Ogura, T. and Hiraga, S. (1986) Structure and Function of the F Plasmid Genes Essential for Partitioning. Journal of Molecular Biology, 192, 1–15.

15. Lin, D.C.-H. and Grossman, A.D. (1998) Identification and Characterization of a Bacterial Chromosome Partitioning Site. Cell, 92, 675–685.

16. Debaugny, R.E., Sanchez, A., Rech, J., Labourdette, D., Dorignac, J., Geniet, F., Palmeri, J., Parmeggiani, A., Boudsocq, F., Anton Leberre, V. et al. (2018) A conserved mechanism drives partition complex assembly on bacterial chromosomes and plasmids. Molecular Systems Biology, 14, e8516.

17. MacCready, J.S., Hakim, P., Young, E.J., Hu, L., Liu, J., Osteryoung, K.W., Vecchiarelli, A.G. and Ducat, D.C. (2018) Protein gradients on the nucleoid position the carbon-fixing organelles of cyanobacteria. eLife.

18. Lim, H.C., Surovtsev, I.V., Beltran, B.G., Huang, F., Bewersdorf, J. and Jacobs-Wagner, C. (2014) Evidence for a DNA-relay mechanism in ParABS-mediated chromosome segregation. eLife, 3.

19. Hwang, L.C., Vecchiarelli, A.G., Han, Y.-W., Mizuuchi, M., Harada, Y., Funnell, B.E. and Mizuuchi, K. (2013) ParA-mediated plasmid partition driven by protein pattern self-organization. The EMBO Journal, 32, 1238–1249.

20. Leonard, T.A., Butler, J. and Löwe, J. (2005) Bacterial chromosome segregation: structure and DNA binding of the Soj dimer — a conserved biological switch. The EMBO Journal, 24, 270–282.

21. Livny, J., Yamaichi, Y. and Waldor, M.K. (2007) Distribution of Centromere-Like parS Sites in Bacteria: Insights from Comparative Genomics. Journal of Bacteriology, 189, 8693–8703.

22. Jensen, R.B. and Gerdes, K. (1997) Partitioning of Plasmid R1. The ParM Protein Exhibits ATPase Activity and Interacts with the Centromere-like ParR-parC Complex. Journal of Molecular Biology, 269, 505–513.

23. Iniesta, A.A. (2014) ParABS System in Chromosome Partitioning in the Bacterium Myxococcus xanthus. PLoS ONE, 9.

24. Barrows, J.M. and Goley, E.D. (2023) Synchronized Swarmers and Sticky Stalks: Caulobacter crescentus as a Model for Bacterial Cell Biology. J Bacteriol, 205, e0038422.

25. Toro, E., Hong, S.H., McAdams, H.H. and Shapiro, L. (2008) Caulobacter requires a dedicated mechanism to initiate chromosome segregation. Proc Natl Acad Sci U S A, 105, 15435–15440.

26. Viollier, P.H., Thanbichler, M., McGrath, P.T., West, L., Meewan, M., McAdams, H.H. and Shapiro, L. (2004) Rapid and sequential movement of individual chromosomal loci to specific subcellular locations during bacterial DNA replication. Proceedings of the National Academy of Sciences, 101, 9257–9262.

27. Bowman, G.R., Comolli, L.R., Zhu, J., Eckart, M., Koenig, M., Downing, K.H., Moerner, W.E., Earnest, T. and Shapiro, L. (2008) A Polymeric Protein Anchors the Chromosomal Origin/ParB Complex at a Bacterial Cell Pole. Cell, 134, 945–955.

28. Ebersbach, G., Briegel, A., Jensen, G.J. and Jacobs-Wagner, C. (2008) A Self-Associating Protein Critical for Chromosome Attachment, Division, and Polar Organization in Caulobacter. Cell, 134, 956–968.

29. Shebelut, C.W., Guberman, J.M., Teeffelen, S.v., Yakhnina, A.A. and Gitai, Z. (2010) Caulobacter chromosome segregation is an ordered multistep process. Proceedings of the National Academy of Sciences, 107, 14194–14198.

30. Dame, R.T., Tark-Dame, M. and Schiessel, H. (2011) A physical approach to segregation and folding of the Caulobacter crescentus genome. Molecular Microbiology, 82, 1311–1315.

31. Jun, S. and Mulder, B. (2006) Entropy-driven spatial organization of highly confined polymers: Lessons for the bacterial chromosome. Proceedings of the National Academy of Sciences, 103, 12388–12393.

32. Kleckner, N., Fisher, J.K., Stouf, M., White, M.A., Bates, D. and Witz, G. (2014) The bacterial nucleoid: nature, dynamics and sister segregation. Current opinion in microbiology, 22, 127– 137.

33. Melendez, A.B., Menikpurage, I.P. and Mera, P.E. (2019) Chromosome Dynamics in Bacteria: Triggering Replication at the Opposite Location and Segregation in the Opposite Direction. mBio, 10.

34. Hester, C.M. and Lutkenhaus, J. (2007) Soj (ParA) DNA binding is mediated by conserved arginines and is essential for plasmid segregation. Proceedings of the National Academy of Sciences, 104, 20326–20331.

35. Schofield, W.B., Lim, H.C. and Jacobs-Wagner, C. (2010) Cell cycle coordination and regulation of bacterial chromosome segregation dynamics by polarly localized proteins. EMBO J, 29, 3068–3081.

36. Thanbichler, M., Iniesta, A.A. and Shapiro, L. (2007) A comprehensive set of plasmids for vanillate- and xylose-inducible gene expression in Caulobacter crescentus. Nucleic Acids Research, 35, e137–e137.

37. Tsai, J.-W. and Alley, M.R.K. (2001) Proteolysis of the Caulobacter McpA Chemoreceptor Is Cell Cycle Regulated by a ClpX-Dependent Pathway. Journal of Bacteriology.

38. Schindelin, J., Arganda-Carreras, I., Frise, E., Kaynig, V., Longair, M., Pietzsch, T., Preibisch, S., Rueden, C., Saalfeld, S., Schmid, B. et al. (2012) Fiji – an Open Source platform for biological image analysis. 9.

39. Lesley, J.A. and Shapiro, L. (2008) SpoT Regulates DnaA Stability and Initiation of DNA Replication in Carbon-Starved Caulobacter crescentus. Journal of Bacteriology, 190, 6867–6880.

40. Menikpurage, I.P., Puentes-Rodriguez, S.G., Elaksher, R.A. and Mera, P.E. (2023) ParA’s Impact beyond Chromosome Segregation in Caulobacter crescentus. J Bacteriol, 205, e0029622.

41. Corrales-Guerrero, L., He, B., Refes, Y., Panis, G., Bange, G., Viollier, P.H., Steinchen, W. and Thanbichler, M. (2020) Molecular architecture of the DNA-binding sites of the P-loop ATPases MipZ and ParA from Caulobacter crescentus. Nucleic Acids Res, 48, 4769–4779.

42. Ptacin, J.L., Lee, S.F., Garner, E.C., Toro, E., Eckart, M., Comolli, L.R., Moerner, W.E. and Shapiro, L. (2010) A spindle-like apparatus guides bacterial chromosome segregation. Nat Cell Biol, 12, 791–798.

43. Hester, C.M. and Lutkenhaus, J. (2007) Soj (ParA) DNA binding is mediated by conserved arginines and is essential for plasmid segregation. Proc Natl Acad Sci U S A, 104, 20326–20331.

44. Collier, J. and Shapiro, L. (2009) Feedback control of DnaA-mediated replication initiation by replisome-associated HdaA protein in Caulobacter. J Bacteriol, 191, 5706–5716.

45. Jensen, R.B., Wang, S.C. and Shapiro, L. (2001) A moving DNA replication factory in Caulobacter crescentus. EMBO J, 20, 4952–4963.

46. Hong, S.H. and McAdams, H.H. (2011) Compaction and transport properties of newly replicated Caulobacter crescentus DNA. Mol Microbiol, 82, 1349–1358.

47. Hong, S.H., Toro, E., Mortensen, K.I., de la Rosa, M.A., Doniach, S., Shapiro, L., Spakowitz, A.J. and McAdams, H.H. (2013) Caulobacter chromosome in vivo configuration matches model predictions for a supercoiled polymer in a cell-like confinement. Proc Natl Acad Sci U S A, 110, 1674–1679.

48. Lindler, L.E., Plano, G.V., Burland, V., Mayhew, G.F. and Blattner, F.R. (1998) Complete DNA sequence and detailed analysis of the Yersinia pestis KIM5 plasmid encoding murine toxin and capsular antigen. Infect Immun, 66, 5731–5742.

49. Jonas, K., Chen, Y.E. and Laub, M.T. (2011) Modularity of the bacterial cell cycle enables independent spatial and temporal control of DNA replication. Curr Biol, 21, 1092–1101.

50. Schwartz, M.A. and Shapiro, L. (2011) An SMC ATPase mutant disrupts chromosome segregation in Caulobacter. Mol Microbiol, 82, 1359–1374.

51. Surovtsev, I.V., Lim, H.C. and Jacobs-Wagner, C. (2016) The Slow Mobility of the ParA Partitioning Protein Underlies Its Steady-State Patterning in Caulobacter. Biophys J, 110, 2790–2799.

52. Shebelut, C.W., Guberman, J.M., van Teeffelen, S., Yakhnina, A.A. and Gitai, Z. (2010) Caulobacter chromosome segregation is an ordered multistep process. Proc Natl Acad Sci U S A, 107, 14194–14198.

53. Bowman, G.R., Comolli, L.R., Gaietta, G.M., Fero, M., Hong, S.H., Jones, Y., Lee, J.H., Downing, K.H., Ellisman, M.H., McAdams, H.H. et al. (2010) Caulobacter PopZ forms a polar subdomain dictating sequential changes in pole composition and function. Mol Microbiol, 76, 173–189.

54. Bowman, G.R., Comolli, L.R., Zhu, J., Eckart, M., Koenig, M., Downing, K.H., Moerner, W.E., Earnest, T. and Shapiro, L. (2008) A polymeric protein anchors the chromosomal origin/ParB complex at a bacterial cell pole. Cell, 134, 945–955.

55. Ebersbach, G., Briegel, A., Jensen, G.J. and Jacobs-Wagner, C. (2008) A self-associating protein critical for chromosome attachment, division, and polar organization in caulobacter. Cell, 134, 956–968.

56. Lasker, K., von Diezmann, L., Zhou, X., Ahrens, D.G., Mann, T.H., Moerner, W.E. and Shapiro, L. (2020) Selective sequestration of signalling proteins in a membraneless organelle reinforces the spatial regulation of asymmetry in Caulobacter crescentus. Nat Microbiol, 5, 418–429.

57. Ptacin, J.L., Gahlmann, A., Bowman, G.R., Perez, A.M., von Diezmann, L., Eckart, M.R., Moerner, W.E. and Shapiro, L. (2014) Bacterial scaffold directs pole-specific centromere segregation. Proc Natl Acad Sci U S A, 111, E2046–2055.

58. Lim, H.C., Surovtsev, I.V., Beltran, B.G., Huang, F., Bewersdorf, J. and Jacobs-Wagner, C. (2014) Evidence for a DNA-relay mechanism in ParABS-mediated chromosome segregation. Elife, 3, e02758.

59. Laloux, G. and Jacobs-Wagner, C. (2013) Spatiotemporal control of PopZ localization through cell cycle-coupled multimerization. J Cell Biol, 201, 827–841.

60. Bowman, G.R., Perez, A.M., Ptacin, J.L., Ighodaro, E., Folta-Stogniew, E., Comolli, L.R. and Shapiro, L. (2013) Oligomerization and higher-order assembly contribute to sub-cellular localization of a bacterial scaffold. Mol Microbiol, 90, 776–795.

61. Murray, H. and Errington, J. (2008) Dynamic control of the DNA replication initiation protein DnaA by Soj/ParA. Cell, 135, 74–84.

62. Kadoya, R., Baek, J.H., Sarker, A. and Chattoraj, D.K. (2011) Participation of chromosome segregation protein ParAI of Vibrio cholerae in chromosome replication. J Bacteriol, 193, 1504–1514.

63. Roberts, D.M., Anchimiuk, A., Kloosterman, T.G., Murray, H., Wu, L.J., Gruber, S. and Errington, J. (2022) Chromosome remodelling by SMC/Condensin in B. subtilis is regulated by monomeric Soj/ParA during growth and sporulation. Proc Natl Acad Sci U S A, 119, e2204042119.

64. Ryter, A. (1965) [Morphologic Study of the Sporulation of Bacillus Subtilis]. Ann Inst Pasteur (Paris), 108, 40–60.

65. Ben-Yehuda, S., Rudner, D.Z. and Losick, R. (2003) RacA, a bacterial protein that anchors chromosomes to the cell poles. Science, 299, 532–536.

66. Wu, L.J. and Errington, J. (2003) RacA and the Soj-Spo0J system combine to effect polar chromosome segregation in sporulating Bacillus subtilis. Mol Microbiol, 49, 1463–1475.

67. Lenarcic, R., Halbedel, S., Visser, L., Shaw, M., Wu, L.J., Errington, J., Marenduzzo, D. and Hamoen, L.W. (2009) Localisation of DivIVA by targeting to negatively curved membranes. EMBO J, 28, 2272–2282.

