## Supplemental Figures for "To let go or not to let go: how ParA can impact the release of the chromosomal anchoring in *Caulobacter crescentus*"

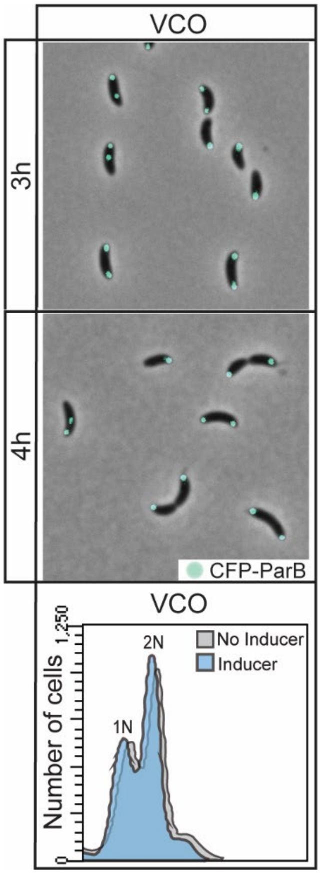


**Supplementary Figure 1.** Wildtype *Caulobacter* cells display only one or two *parS* loci A mixed population of CB15N cells expressing *parB::cfp-parB* (cyan) were diluted to 0.1 $OD_{600,}$ then added 0.1% xylose to express *xylX::empty-vector*; then micrographs were obtained at **A)** 3 h or **B)** 4 h of induction. Micrograph scale bar represents 2 μM. **C)** Flow cytometry profiles after 3 h of expression. Flow cytometry data were obtained by synchronizing the aforementioned cells and adding 0.1% xylose for 3 h, then adding Rifampicin to block re-initiation of chromosome replication for an additional 3 h. Chromosomal content is denoted above each peak.

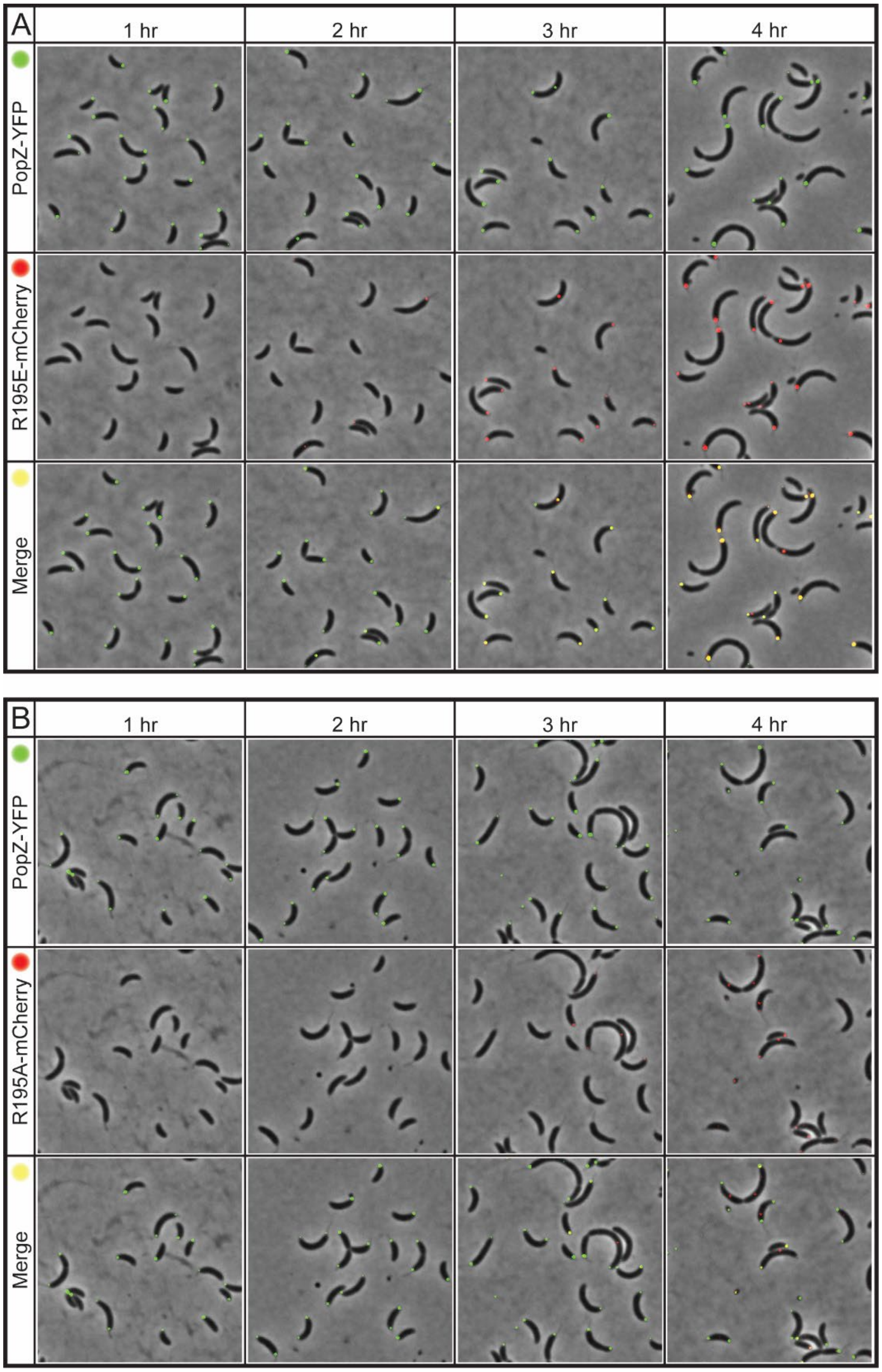


**Supplementary Figure 2.** Correlation between ParA-mCherry appearance post induction and PopZ-YFP localization. Micrographs of CB15N cells expressing *popZ::popZ-yfp* and **A)** *xylX::parA-(R195E)-mCherry* or **B)** *xylX::parA-(R195A)-mCherry*. Images were taken at 1 h intervals until 4 h to observe the localization of ParA and PopZ.

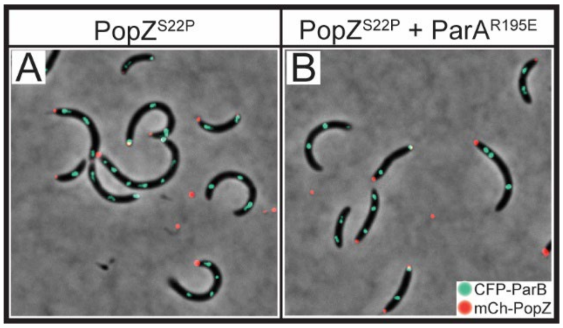


**Supplementary Figure 3.** Expression of *popZ*-S22P with wildtype ParA results in replication and segregation defects. Micrographs of CB15N cells expressing *parB::cfp-parB* and **A)** *popZ::mCherry-popZ-(S22P)* or **B)** *popZ::mCherry-popZ-(S22P)* with *xylX::parA-(R195E).* Micrographs are representative of the cell population.
